## Supplementary material for "Low breeding propensity and success of European Turtle Doves in Germany"

#### Contents:

1. **Short summary of nest site identification**
2. **Tables S1-S4**
3. **Fig. S1**
4. **Full workflow for the breeding analysis**

#### Short summary of nest site identification

##### Identification of potential nest sites

To identify potential nest sites, we searched for GPS data clusters of an individual in the QGIS visualization. The "Points to Path" tool was used to determine whether a cluster had been visited on at least 13 consecutive days. The center of each of these potential nest sites was buffered by 45 meters (= potential breeding area) to account for scattering of the GPS data due to terrain type, tree cover, etc.. The tracking data were intersected with the potential breeding area to create hourly summaries of the presence in the breeding area. Presence per hour was calculated as a percentage of the positions in the nesting area, compared with the home range (95% kernel) data. For data gaps, we assumed that the average presence during these gaps matched the actual presence.

##### Confirmation of potential nest sites via nestR

When utilizing nestR we used very loose conditions (Table 1), as recommended by Picardi (2020). The result was a list of 93 potential nest sites. Comparing the coordinates to our own findings (Table 2), we detected that the first entry for each individual matched our own results differing with distances ranging from 6 – 38 m (Ø 18.8 m ± 9.9 m). Distances were calculated in R using the distGeo function from the geosphere package (Karney 2013). As the distances are well within the chosen 45 m buffer size, we would recommend further studies using nestR directly to detect potential nest sites by simply taking the first entry/entries for each individual.

Reference: Karney, C. F. (2013). Algorithms for geodesics. *Journal of Geodesy*, *87*, 43-55.

***Table S1**: Settings for the nestR: find_nests function to determine the nest coordinates of the GPS-tagged European Turtle Doves.*

| Parameter explanation, copied from the nestR helpfunktion in R | Parameter | our setting |
|---|---|---|
| data.frame of movement data. Needs to include burst, date, long, lat | gps_data | |
| Character string. Earliest date to be considered within the breeding season. Month and day, format "mm-dd" | sea_start | 05-01 |
| Character string. Latest date to be considered within the breeding season. Month and day, format "mm-dd" | sea_end | 08-31 |
| Duration of nesting cycle | nest_cycle | 31 |
| Size of the buffer to compute location revisitation | buffer | 45 |
| Minimum number of points within a buffer | min_pts | 2 |
| Minimum number of fixes for a day to be retained if no nest visit was recorded | min_d_fix | 5 |
| Minimum number of consecutive days visited | min_consec | 13 |
| Minimum percent of fixes at a location on the day with maximum attendance | min_top_att | 1 |
| Minimum percent of days spent at a location between first and last visit | min_days_att | 1 |
| If results include temporally overlapping attempts, select only one among those? Defaults to TRUE. | discard_overlapping | FALSE |

***Table S2****: Comparison of nest coordinates for European Turtle Doves obtained with nestR to the study coordinates using distGeo (package geosphere)*

| **ID** | **No. Incubation** | **Position nestR list** | **Long** | **Lat** | **NestR_long** | **NestR_lat** | **Distance [m]** |
|---|---|---|---|---|---|---|---|
| TT-23-HE-01 | 1 | 1 | 8.8696 | 50.8619 | 8.8697 | 50.8618 | 10 |
| TT-23-HE-02 | 1 | 1 | 8.8696 | 50.8619 | 8.8697 | 50.8620 | 9 |
| TT-23-HE-03 | 1 | 1 | 8.7993 | 51.2718 | 8.7993 | 51.2719 | 12 |
| TT-23-HE-06 | 1 | 1 | 8.2649 | 50.4540 | 8.2647 | 50.4540 | 17 |
| TT-23-SA-01 | 1 | 1 | 12.2138 | 51.1593 | 12.2137 | 51.1594 | 18 |
| TT-24-BB-04A | 1 | 1 | 14.4884 | 52.0189 | 14.4882 | 52.0190 | 17 |
| TT-24-BB-04B | 1 | 1 | 13.0349 | 51.6179 | 13.0349 | 51.6180 | 9 |
| TT-24-HE-01 | 1 | 1 | 9.0595 | 50.3472 | 9.0597 | 50.3475 | 38 |
| TT-24-HE-03 | 2 | 1 | 8.8856 | 49.9779 | 8.8859 | 49.9779 | 18 |
| TT-24-HE-03 | 1 | 2 | 8.8874 | 49.9802 | 8.8873 | 49.9801 | 21 |
| TT-24-HE-04 | 2 | 1 | 9.0546 | 50.3477 | 9.0543 | 50.3479 | 32 |
| TT-24-HE-04 | 1 | 2 | 9.0558 | 50.3474 | 9.0556 | 50.3476 | 33 |
| TT-24-HE-05 | 1 | 1 | 8.2985 | 50.4398 | 8.2985 | 50.4399 | 6 |
| TT-24-HE-05 | 2 | 2 | 8.3011 | 50.4403 | 8.3012 | 50.4403 | 8 |
| TT-24-HE-06 | 1 | 1 | 9.0538 | 50.3490 | 9.0536 | 50.3492 | 21 |
| TT-24-SA-01 | 1 | 1 | 12.1948 | 51.1305 | 12.1947 | 51.1305 | 7 |
| TT-24-SA-02 | 2 | 1 | 12.2100 | 51.1502 | 12.2102 | 51.1505 | 33 |
| TT-24-SA-02 | 1 | 2 | 12.2089 | 51.1487 | 12.2090 | 51.1488 | 12 |
| TT-24-SA-06 | 1 | 1 | 12.1953 | 51.1346 | 12.1953 | 51.1347 | 10 |
| TT-24-TH-03 | 1 | 1 | 11.2730 | 50.8232 | 11.2730 | 50.8234 | 17 |
| TT-24-TH-03 | 2 | 2 | 11.2745 | 50.8246 | 11.2742 | 50.8244 | 26 |
| TT-24-TH-06 | 1 | 1 | 10.5264 | 51.0945 | 10.5267 | 51.0945 | 23 |
| TT-24-TH-07 | 1 | 1 | 10.5264 | 51.0945 | 10.5268 | 51.0943 | 36 |

***Table S3:*** *European Turtle Doves (Streptopelia turtur) equipped with GPS-GSM transmitters (n = 45).*

| ID | Season | Federal state | Transmitter ID | Deployment date [dd mm yyyy] | Capture location DE [lat, long] |
|---|---|---|---|---|---|
| TT-23-HE-01 | 2023 | Hesse | 0865 | 02.06.2023 | 50.863; 8.867 |
| TT-23-HE-02 | 2023 | Hesse | 07E0 | 15.06.2023 | 50.863; 8.867 |
| TT-23-HE-03 | 2023 | Hesse | 07CF | 20.06.2023 | 51.269; 8.793 |
| TT-23-HE-04 | 2023 | Hesse | 07E7 | 20.06.2023 | 51.269; 8.793 |
| TT-23-HE-05 | 2023 | Hesse | 6717 | 29.06.2023 | 50.457; 8.225 |
| TT-23-HE-06 | 2023 | Hesse | 6718 | 05.07.2023 | 50.445; 8.256 |
| TT-23-SA-01 | 2023 | Saxony-Anhalt | 07C4 | 06.06.2023 | 51.144; 12.146 |
| TT-23-SA-02 | 2023 | Saxony-Anhalt | 0802 | 07.06.2023 | 51.144; 12.146 |
| TT-23-SA-03[a] | 2023 | Saxony-Anhalt | 07C3 | 07.06.2023 | 51.144; 12.146 |
| TT-23-SA-04[a] | 2023 | Saxony-Anhalt | 07D4 | 07.06.2023 | 51.144; 12.146 |
| TT-23-SA-05 | 2023 | Saxony-Anhalt | 07A6 | 08.06.2023 | 51.144; 12.146 |
| TT-23-SA-06[a] | 2023 | Saxony-Anhalt | 07CC | 24.06.2023 | 51.144; 12.146 |
| TT-24-BB-01 | 2024 | Brandenburg | OA 00 | 20.06.2024 | 52.001; 14.475 |
| TT-24-BB-02[b] | 2024 | Brandenburg | 244893 | 21.06.2024 | 52.001; 14.475 |
| TT-24-BB-03[b] | 2024 | Brandenburg | 244895 | 21.06.2024 | 52.001; 14.475 |
| TT-24-BB-04 | 2024 | Brandenburg | OA 74 | 21.06.2024 | 52.001; 14.475 |
| TT-24-BB-05[c] | 2024 | Brandenburg | 54226 | 22.06.2024 | 52.001; 14.475 |
| TT-24-HE-01 | 2024 | Hesse | 243973 | 30.05.2024 | 50.348; 9.055 |
| TT-24-HE-02[a] | 2024 | Hesse | 243970 | 30.05.2024 | 50.348; 9.055 |
| TT-24-HE-03 | 2024 | Hesse | 243978 | 31.05.2024 | 49.929; 8.857 |
| TT-24-HE-04 | 2024 | Hesse | 243974 | 01.06.2024 | 50.348; 9.055 |
| TT-24-HE-05 | 2024 | Hesse | 243968 | 03.06.2024 | 50.445; 8.259 |
| TT-24-HE-06 | 2024 | Hesse | 243969 | 06.06.2024 | 50.349; 9.055 |
| TT-24-HE-07 | 2024 | Hesse | 130000124E | 08.06.2024 | 50.445; 8.256 |
| TT-24-HE-08 | 2024 | Hesse | 1300001200 | 08.06.2024 | 50.445; 8.256 |
| TT-24-HE-09 | 2024 | Hesse | 130000121A | 08.06.2024 | 50.445; 8.256 |
| TT-24-HE-10[d] | 2024 | Hesse | 1300000977 | 28.06.2024 | 51.272; 8.873 |
| TT-24-HE-13 | 2024 | Hesse | 241580 | 17.07.2024 | 51.272; 8.873 |
| TT-24-SA-01 | 2024 | Saxony-Anhalt | 8964 | 17.06.2024 | 51.144; 12.146 |
| TT-24-SA-02 | 2024 | Saxony-Anhalt | 8968 | 18.06.2024 | 51.144; 12.146 |
| TT-24-SA-04 | 2024 | Saxony-Anhalt | 8966 | 20.06.2024 | 51.144; 12.146 |
| TT-24-SA-05[a] | 2024 | Saxony-Anhalt | 6719 | 20.06.2024 | 51.144; 12.146 |
| TT-24-SA-06 | 2024 | Saxony-Anhalt | 8965 | 20.06.2024 | 51.144; 12.146 |
| TT-24-TH-01 | 2024 | Thuringia | 243971 | 05.06.2024 | 50.795; 11.263 |
| TT-24-TH-02 | 2024 | Thuringia | 243972 | 06.06.2024 | 50.795; 11.263 |
| TT-24-TH-03 | 2024 | Thuringia | 243975 | 06.06.2024 | 50.795; 11.263 |
| TT-24-TH-04[b] | 2025 | Thuringia | 244892 | 10.06.2024 | 50.795; 11.263 |
| TT-24-TH-05[b] | 2026 | Thuringia | 244890 | 10.06.2024 | 50.795; 11.263 |
| TT-24-TH-06 | 2027 | Thuringia | 243976 | 12.06.2024 | 51.095; 10.533 |
| TT-24-TH-07 | 2028 | Thuringia | 243977 | 12.06.2024 | 51.095; 10.533 |
| TT-24-TH-08 | 2029 | Thuringia | 243979 | 13.06.2024 | 51.037; 10.458 |
| TT-24-TH-09[b] | 2030 | Thuringia | 244894 | 14.06.2024 | 51.095; 10.533 |
| TT-24-TH-10[b] | 2031 | Thuringia | 244896 | 17.06.2024 | 50.795; 11.263 |
| TT-24-TH-11 | 2032 | Thuringia | 1300001208 | 17.06.2024 | 50.795; 11.263 |
| TT-24-TH-12 | 2033 | Thuringia | 8969 | 17.06.2024 | 50.795; 11.269 |

[a] too little GPS data, [b] no GPS data, [c] no animal movements, [d] transmitter removed (02.07.2024)

***Table S4:*** *R settings (grid, extent) for the kernel density estimation of the European Turtle Doves (n = 32) using Epanechnikov kernels (kern = epa). The smoothing parameter (h) was set to: href for all individuals. For individuals with breeding season displacement (TT-24-BB-04, TT-24-TH-08) the letters A and B were added to the ID to indicate the different locations.*

| **ID** | **grid** | **extent** |
|---|---|---|
| TT-HE-23-01 | 200 | 0.5 |
| TT-HE-23-02 | 200 | 0.5 |
| TT-HE-23-03 | 200 | 0.5 |
| TT-HE-23-04 | 200 | 0.5 |
| TT-HE-23-05 | 200 | 0.5 |
| TT-HE-23-06 | 200 | 0.5 |
| TT-SA-23-01 | 200 | 0.5 |
| TT-SA-23-02 | 200 | 0.5 |
| TT-SA-23-05 | 200 | 0.5 |
| TT-24-BB-01 | 100 | 1.0 |
| TT-24-BB-04A | 200 | 1.0 |
| TT-24-BB-04B | 100 | 1.0 |
| TT-24-HE-01 | 200 | 1.0 |
| TT-24-HE-03 | 100 | 1.0 |
| TT-24-HE-04 | 150 | 1.0 |
| TT-24-HE-05 | 200 | 1.0 |
| TT-24-HE-06 | 200 | 1.0 |
| TT-24-HE-07 | 200 | 1.0 |
| TT-24-HE-08 | 150 | 1.0 |
| TT-24-HE-09 | 100 | 1.0 |
| TT-24-HE-13 | 150 | 0.5 |
| TT-24-SA-01 | 200 | 2.0 |
| TT-24-SA-02 | 300 | 1.0 |
| TT-24-SA-04 | 100 | 1.0 |
| TT-24-SA-06 | 200 | 2.0 |
| TT-24-TH-01 | 200 | 3.0 |
| TT-24-TH-02 | 200 | 1.0 |
| TT-24-TH-03 | 200 | 1.0 |
| TT-24-TH-06 | 200 | 3.0 |
| TT-24-TH-07 | 150 | 1.0 |
| TT-24-TH-08A | 200 | 6.0 |
| TT-24-TH-08B | 150 | 1.0 |
| TT-24-TH-11 | 200 | 3.0 |
| TT-24-TH-12 | 150 | 1.0 |

**Fig. 1** Capture locations of European Turtle Doves for GPS tracking (2023 – 2024) in four federal states of Germany.

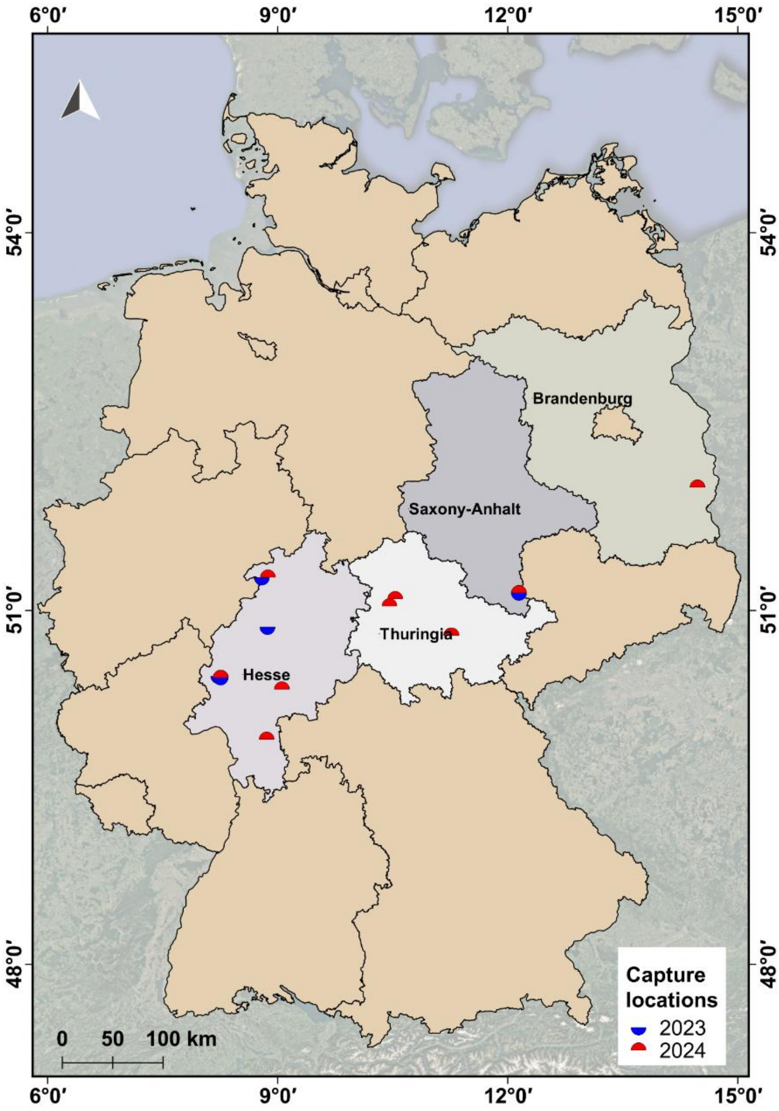

### Workflow for suspected breeding analysis – European Turtle Dove

#### 1. Preparing dove data in Excel

##### 1.1 Preparing the data

The raw data should remain unaltered. Therefore, after opening the file, immediately save it under a new name. Delete unnecessary columns. For habitat analysis, you need the columns with the dove ID, both coordinate columns, and the timestamp.

Important: Discuss with the project lead in advance which timestamp should be used. In the TT data, for example, there are columns labeled 'timestamp' and 'study-local-timestamp', with time entries in Central European Summer Time (CEST), which should be used for our analyses!

Arrange the columns as follows: ID as the first column, followed by the coordinate columns, then 'timestamp' or 'study-local-timestamp'. Rename the coordinate columns to 'lon' and 'lat'.

##### 1.2 Splitting the chosen timestamp column

Label the chosen timestamp column as 'timestamp', copy it, and paste it as a new column (E).

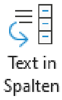


Go to the 'Data' tab. Use the 'Text to Columns' function. The Text Conversion Assistant will open. Continue by left-clicking. Under delimiter options, select 'Space', then click 'Finish'. Rename the second timestamp column to 'date'. Right-click on the 'date' column, then left-click 'Format Cells', go to the 'Number' tab, select 'Date', and under locale choose 'German', then select the line with format: 'YYYY-MM-DD'. Confirm with OK.

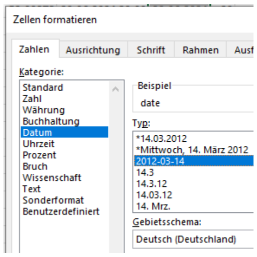


Label the new column as 'time' and move it so that three empty columns follow behind 'date'. Copy the 'date' column and paste it behind itself. Select the second 'date' column, go to the 'Data' tab, and use 'Text to Columns' again. Under delimiters, select 'Other' and enter a dot (.), then click 'Finish'. Rename the resulting columns to: 'day' (F), 'month' (G), 'year' (H). Format 'day' as a number with 0 decimal places.

Copy the 'time' column and paste it behind itself. Select the second 'time' column, go to the 'Data' tab, and use 'Text to Columns'. Under delimiters, choose 'Other' and enter a colon (:), then click 'Finish'. Rename the resulting columns to: 'hour' (J), 'minute' (K), and delete the last column (seconds). Format 'hour' as a number with 0 decimal places.

Right-click on the column 'hour', then left-click on 'Format Cells'. Click on the 'Number' tab. Under Category, left-click on 'Number'. Set the decimal places to 0 and finally, left-click on 'OK'. Result:

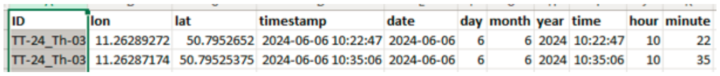

| ID | lon | lat | timestamp | date | day | month | year | time | hour | minute |
|---|---|---|---|---|---|---|---|---|---|---|
| TT-24_Th-03 | 11.26289272 | 50.7952652 | 2024-06-06 10:22:47 | 2024-06-06 | 6 | 6 | 2024 | 10:22:47 | 10 | 22 |
| TT-24_Th-03 | 11.26287174 | 50.79525375 | 2024-06-06 10:35:06 | 2024-06-06 | 6 | 6 | 2024 | 10:35:06 | 10 | 35 |

##### 1.3 Create Excel files per dove

Select the 'Home' tab. Use a left-click to activate the 'ID' column. Right-click on 'Filter' in the ribbon under 'Sort & Filter'.

Left-click on the dropdown arrow that appears in the 'ID' column. All doves contained in the Excel file will be listed there. Untick 'Select All',left-click on the first individual and left-click on 'OK'.

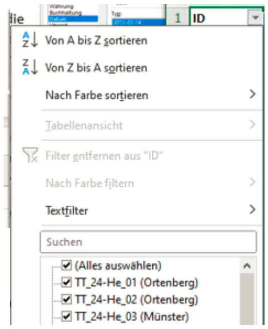

Open a new blank Excel workbook. Copy the header row from the 'Master Workbook' and paste it into row 1 of the blank workbook. Save the new file as: CSV (Comma delimited) (*.csv) at your desired location with the dove ID as the file name.

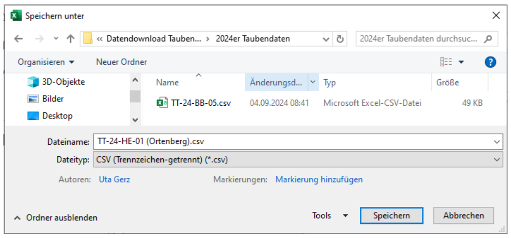


Click in a data cell of the master workbook and use CTRL+A to select all rows with content, then copy them with CTRL+C. In the new .csv file, click in cell A2 and use 'Paste Special' to insert the copied content there.

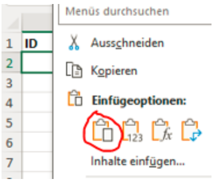


##### 1.4 Clean data

Identify the data that precedes the capture time and delete the corresponding rows. If the start time of the individual's migration is already known, delete the subsequent data as well. Save the file. This is the data preparation for QGIS.

##### 1.5 Create an Excel workbook for each individual

To ensure all data for the individual is organized clearly later, create an .xlsx file. Open the .csv file of the individual's GPS data, save it as an .xlsx file, and leave it open. Name the worksheet as GPS_ID (of the individual).

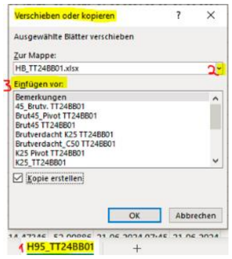


Add future .csv data as worksheets. To do this, open the relevant (.csv) file, left-click on the tab with the worksheet name (1), and in the window that opens, select the Move or Copy option with a left-click. In the next window that opens, left-click on the dropdown arrow to the right (2) and select the individual's .xlsx file. You can choose where in the workbook the worksheet should be placed (3), then confirm with OK. Note: If you do not tick the box for Create a Copy, this worksheet will typically not be available in the original workbook anymore. Save and close the original workbook.

##### 1.6 Data preparation for R

Save the resulting .csv as a .txt file (Tab Delimited) (*.txt) at your desired location. Note: Hyphens should not be used in this filename, as R might interpret them as minus signs.

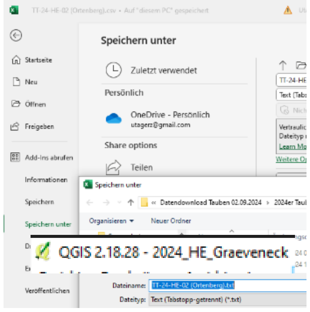


Retain only the first three columns (ID, lon, lat), and delete the remaining columns. Save the file. Repeat the process for the other individuals. By now, there should be a .csv file, a .txt file, and an .xlsx workbook for each individual. Check for completeness!

#### 2. Work in QGIS (version 2.18.28 Las Palmas)

Open a project in QGIS, give it a clear name (possibly including the year and location), and save it at your desired location. If dove data from multiple individuals exist at a single capture location, it is advisable to include as many of these individuals in the project as possible to easily identify overlaps in habitats and similar patterns at a glance.

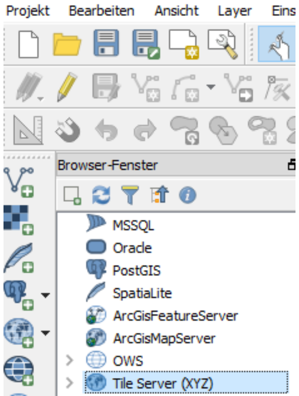


##### 2.1 Start the project

In the Browser window (left), right-click on 'Tile Server (XYZ)', and the option 'New Connection' will appear. In the window that opens: 'New XYZ Tile Layer', paste this address: **https://mt1.google.com/vt/lyrs=s&x={x}&y={y}&z={z}** and

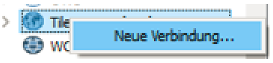


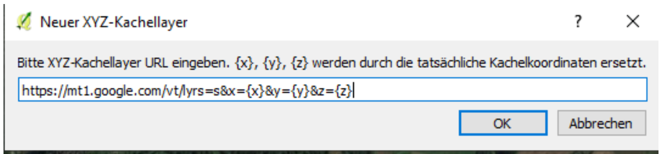

confirm with a left-click on 'OK'. A further window will open where you need to enter a name, e.g., 'Google Satellite'. After entering, confirm with an left-click on 'OK'. Note: When publishing maps created with this method, pay attention to Google's data usage terms and license rights!

Now the map is available as a layer. If the new connection is not immediately visible, left-click on the greater-than sign next to the Tile Server. This will expand the available options. With a double left-click on the appropriate tile, the Google Satellite layer will appear in the layer window below. It contains just the 'bare' landscape without place names. This can be disadvantageous because, he place labels are missing. Therefore, additionally load the Google Hybrid map: **https://mt1.google.com/vt/lyrs=y&x={x}&y={y}&z={z}.** When the Google Hybrid layer is activated with a left-click in the layer window, a checkmark appears in front of the layer, and roads, place names, etc., will appear on the map. However, they can often be distracting, for instance, in print compositions.

##### 2.2 Districts, parcels, land plots (optional)

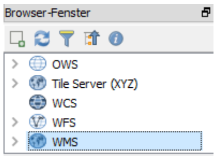


To display districts, parcels, and land plots later if needed, right-click in the browser panel on: WMS, and select New Connection.

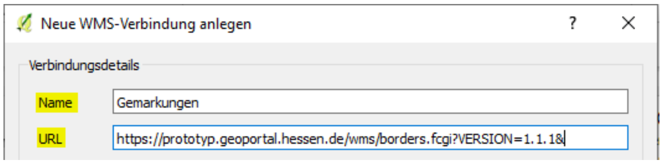


A window opens where you can enter a name, e.g., Districts, and input the following URL for the state of Hesse: https://prototyp.geoportal.hessen.de/wms/borders.fcgi?VERSION=1.1.1&.

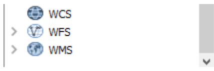


Left-Click OK. The data is now available as a layer. If the new connection is not visible right away, click the arrow (greater-than sign) to the left of WMS. The available options will expand.

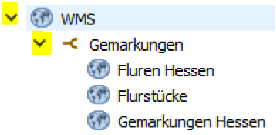


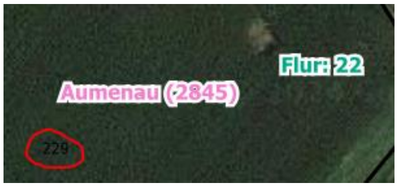


Click the arrow next to Districts (or Parcels**,** Plots**,** Districts Hesse). Double-left-click on the relevant entries to display them as individual layers in the layer panel. Note: The data from active layers will only become visible at a scale of 1:5000 or smaller. Labels may overlap on the map, so activate/deactivate them individually as needed.

##### 2.3 Load GPS data of an individual

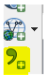

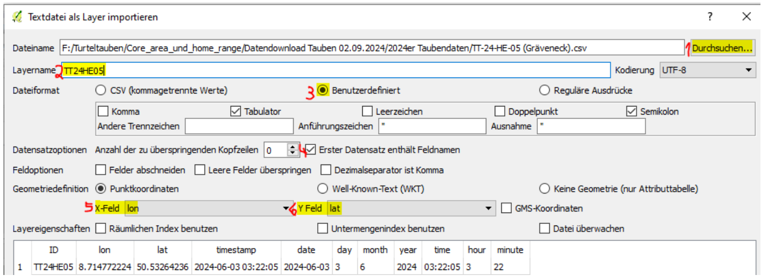


Left-click on the comma icon, a window will open. Click on Browse (1) and select the desired .csv file. Complete all remaining inputs (2-6) if they are not automatically filled in correctly. In the layer panel, the new layer will automatically appear using coordinate reference system (CRS) 4326, and the GPS data will be displayed on the map as points.

##### 2.4 Change CRS of the point layer

To accurately calculate the sizes of the kernels (home range, core area, etc.) later, their data must be in a projected CRS (for Germany: CRS 4647). The point layer data will later be intersected with the polygon layers of the kernels. Both layers must be in the same CRS for this.Therefore, the generated point layer has to be saved immediately again using CRS 4647:

Right-click on the layer → choose Save as… (A)

In the new window:

1. Select ESRI Shapefile as the format from the dropdown menu.
2. Click Browse to specify a custom save location and file name.
3. You can reuse the same name as before.
4. Under CRS, select EPSG:4647 from the dropdown menu.
5. Make sure to tick the box: Add saved file to map so that the layer is automatically added to the project.

Click OK to finish. The new layer will appear on the map.

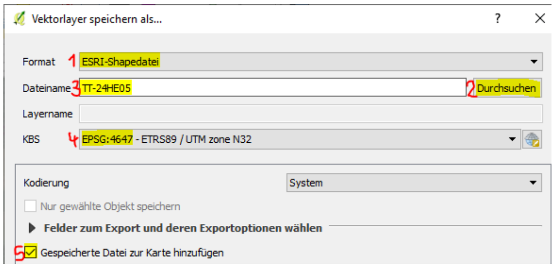


To improve clarity, remove the original (import) layer: Click the original layer in the layer panel. In the opened menu, click Remove (B) and confirm.

Additional options in this menu include: Open attribute table (C) and edit layer properties — e.g., under Style (D), change the symbol, size, or color of the points.

Tip: Use clearly distinguishable colors for each individual that contrast well with the background

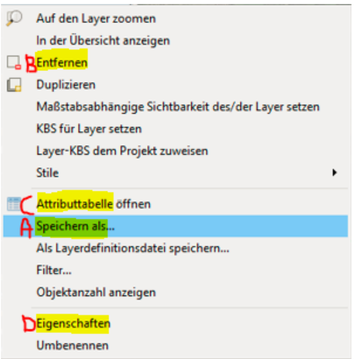


#### 2.5 Daily routes with the Point-to-Path Tool

To visualize daily routes/paths: Open the Processing Toolbox via the Processing tab. It will appear on the right side of the screen.

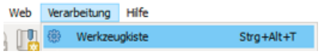


Use the Search function to find Points to Path and double-left-click to open it.

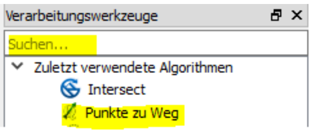


In the opened window, for Input point layer, select the GPS point layer of the individual in the dropdown menu. Ensure that the box for Open output file after running algorithm is ticked. For both Group field and Order field, select date from the dropdown menu. Click Run. → The paths will now appear on the map, and a new layer named Paths will be added to the layer panel. To ensure the paths are saved and assigned to the individual: Click on the Paths layer. Follow the same steps as described in section 2.4 (CRS change and saving). Save the layer under a clear name, e.g., Routes_TT24HE05.To improve clarity: Right-click the original temporary path layer. Select Remove and confirm removal.

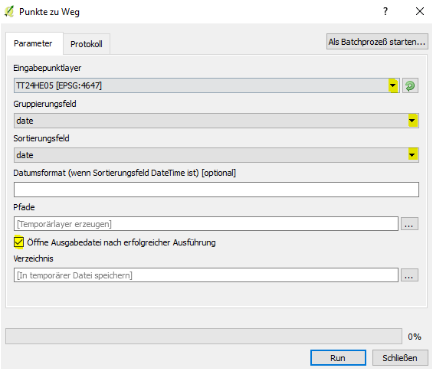


#### 2.5 Categorize routes

To better identify the daily routes, they are categorized: Right-click the path layer and click on the line: Properties. A window opens. On the left, click the tab: Style. In the top dropdown menu, select: Categorized. In the dropdown for column: choose abc group. For color scheme: select Random Colors. Below the (initially empty) field, click on:

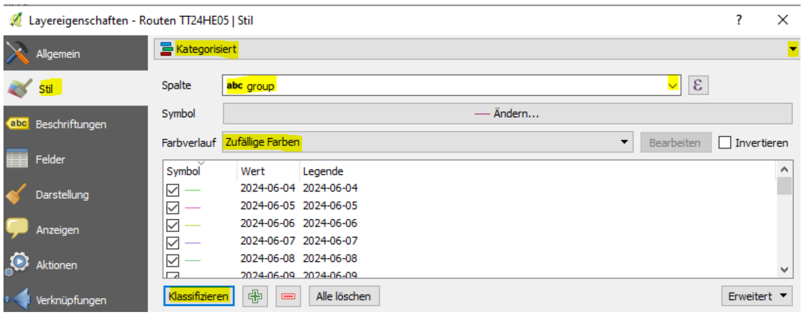

Classify. The field now fills with entries. Scroll to the bottom, select the last entry and delete it using the minus symbol next to Classify. Click OK. → The paths on the map now appear in color. The symbol for the routes in the layer panel has changed.

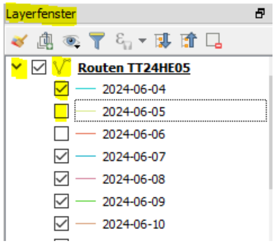


Next to the layer is a dropdown arrow. If pointing downward, all entries are visible and can be (de)selected individually. This makes it easier to view routes from specific time periods and recognize patterns. Repeated, similar routes over extended periods may indicate a breeding attempt. Highly variable, wide-ranging routes suggest exploratory behavior. You can right-click any date entry to show/hide all days at once, instead of (un)ticking each one manually.

Note: Some individuals may undergo a secondary migration to a more distant area (e.g., if they do not find a partner or if the breeding attempt fails). This is especially visible in route analysis. In such cases, consider splitting the GPS data and analyzing each area/timeframe separately. For kernel calculation, you can rename these with identifiers like ID_A.

#### 3. Kernel calculation in R – Searching for a 'slim fit'

The kernel calculation (home range, core area, etc.) is done here with the package adehabitatHR. The goal is to obtain kernels that have little empty space around the GPS data. Information on home range and core area calculation can be found at: http://scbi-migbirds.github.io/homeRange.html

##### 3.1 Preparation in R

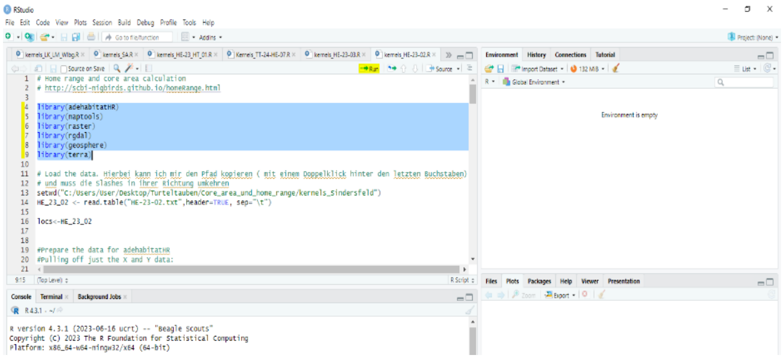

Open RStudio. The four main panels are displayed (information copied from Ecosia AI Chat).

Source Pane (top left): Here you can create and edit R scripts, markdown files, or other code files.
Console Pane (bottom left): This is the area where you can directly enter and execute R commands. You will also see the outputs of your commands here.
Environment/History Pane top right): This area displays the current objects and variables you have created in your session. You can also find the history list of previously entered commands.
Files/Plots/Packages/Help Pane (bottom right): This area is multifunctional. Here you can browse your files, display plots, manage packages, and access help.

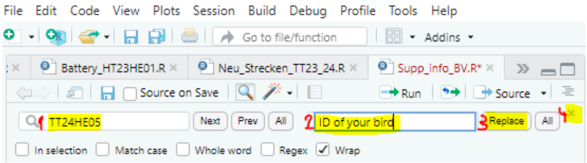


Up to this point, the procedure is the same for all individuals. In the following part, you need to experiment with the settings to find which ones work and produce good results. Good results are characterized by the smallest possible kernels around the GPS data. However, there must still be enough data within the kernel. This can often only be determined through further work in QGIS.

The following information is copied from Ecosia AI Chat.
- kud is the object in which the result of the kernel density estimation is stored.
- kernelUD() is the function that calculates the kernel density estimation.
- h = 'href': This specifies the method for estimating the bandwidth (h). 'href' stands for 'href method', a method for estimating the optimal bandwidth for the kernel density estimation.
- locs1[, 1] refers to the first column of the data frame or matrix locs1, which contains the location data. The use of [, 1] means that only the values from the first column are selected.
- grid specifies the number of grid points used for the density calculation. A higher value can lead to a more detailed density estimation.
- extent specifies the extent over which the density estimation is carried out. A value of 3 means, for example, that the density estimation is conducted over a range of three units around the data points.
- kern = 'epa': This means that the Epanechnikov kernel is used.

##### 3.4 Visualization in R

Drawing the vector and the points in the plotting window under the 'Plots' tab will display a graphic with the GPS data and the calculated kernels. The H95 is outlined in green, C50 in blue, and K25 in red. Often, only H95 and C50 are necessary. For our data from TT24HE05, the displayed result was obtained. It looks good in R, but less so in QGIS, therefore we continued searching by varying the settings of grid and extent. However, in this case it was the best obtained alternative…

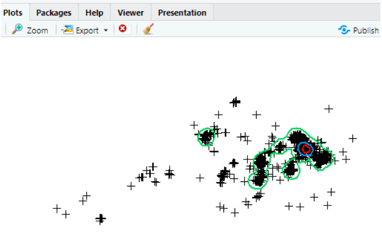


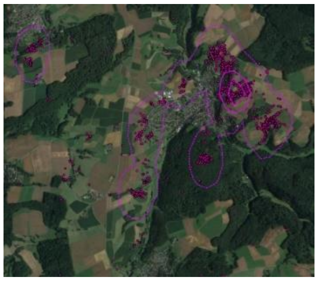

Copy (into the source pane) and Run lines 64-69 of the R-Script: **Supp_info_BV.R.**, modify them to match your individual, and then run all lines with a left mouse click on Run.

##### 3.5 Create kernel shapefiles

The created shapefiles will be saved in the directory specified in the 'Load Data in R' section. In the example: storage at F:/Turtledoves/Core_area_and_home_range/DataDownload_2024-09-02/2024_Turtledove_Data.

Copy (into the source pane) and Run lines 71-75 of the R-Script: **Supp_info_BV.R.**, modify them to match your individual, and then run all lines with a left mouse click on Run.

##### 3.6 Import kernel polygons as a layer in QGIS

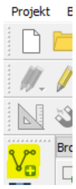


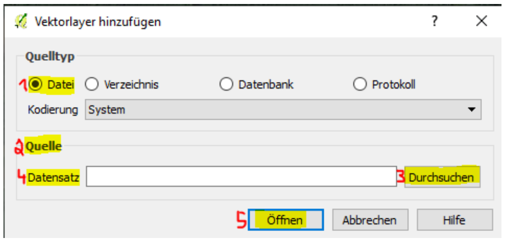


In the open QGIS project, the polyscapes stored in the desired directory are added as layers by selecting the 'Add Vector Layer' function. A window opens, and you keep the ticked setting for Source Type as File (1) and choose the location under Source (2) by left-clicking on Browse (3). In the opened window of the storage location, select the .sph file of the polyshapes you want to import (HR95 or C50...) with a double-left-click, which fills the Dataset (4) line. By left-clicking on Open (5), the polyscape is imported as a layer in CRS 4326 and appears in the layer list and on the map.

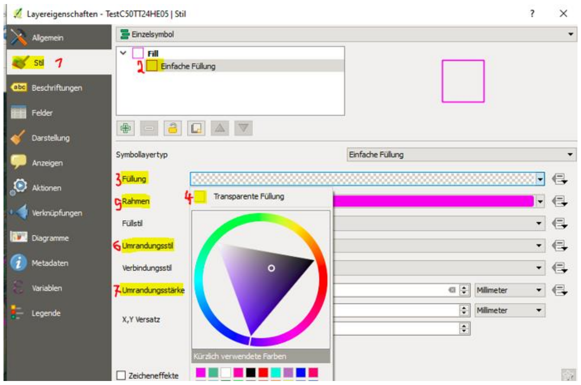

However, for later area calculation and intersection with GPS data, CRS: 4647 is required, so proceed analogously to 2.4. The area displayed on the map is filled. This is changed by opening the Layer Properties of the layer. Click on the layer in the layer panel, selecting Style (1), and clicking on Simple Fill (2). In the modified window, click on the drop-down menu of the Fill (3) line. Another small window opens. Click on Transparent Fill (4). Click on the line: Stroke (5), and set the stroke color there. For better clarity with multiple individuals on one map, you can choose the same stroke color for all kernel shapes of an individual and vary the border style. For border style (6), choose dotted line for HR95, solid line for C50, and dashed line for K25. For border width (7), choose 0.66. Confirm the changed settings by clicking OK.

##### 3.7 Intersection of GPS point layer data with kernel polygons

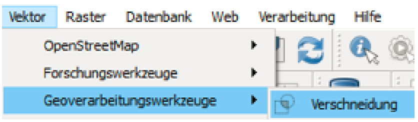


To determine the point data of the individual within a kernel, they are intersected. The intersection data is needed to determine nesting suspicion. Under the 'Vector' tab, select left-clicking the line: 'Geoprocessing Tools' and there: 'Intersection'. The intersection window opens. Left-click on the drop-down menu of the input layer, then left-click on the point layer of the individual. Left-click on the drop-down menu of: 'Intersect layer' and left-click on the kernel to be intersected. Tick the boxes 'Ignore null geometries' and 'Open output file after running' and with a left-click on: 'Run', the process starts. A (temporary) layer named 'Intersection' appears in the layer panel and the associated points on the map. Since every intersection layer generated by QGIS is named identically, it is advisable to edit it further immediately if one wants to keep the layer. Alternatives include saving as a .csv file or using the 'Save As... ' function to create a permanent layer (optional). Usually, saving as a .csv file is sufficient.

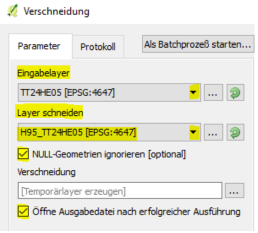

##### 3.8 Save intersection data as .csv

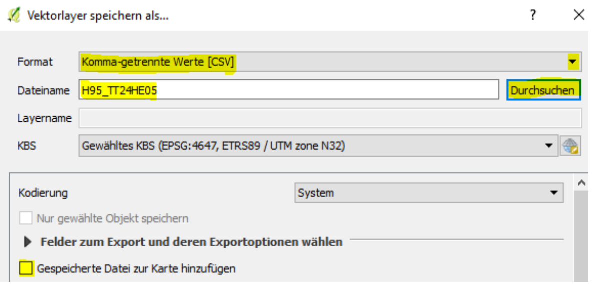


The intersection data of the kernels is needed as .csv files for further work. In the layer panel, left-click on the resulting intersection layer and right-click select 'Save As'. In the window that opens: 'Save Vector Layer As...', left-click on the drop-down menu of the 'Format' line, and choose the option: 'Comma Separated Values [CSV]'. Left-click on 'Browse', set the storage location, and then enter a name under 'File name'. Since the file is not needed as a layer, remove the default checkmark by unticking the box before 'Add saved file to map', and then left-click 'OK' at the bottom. To determine nesting suspicion, the .csvs of at least H95 is required. After successful export, you can delete the intersection layer from the layer panel by right-clicking it and then selecting 'Remove', confirming to delete it as usual.

#### 4. Nesting suspicion (determine for core area) – alternatively use nestR directly (4.9)

We found that the R package: nestR (Picardi et al. 2020) could be used effectively, but achieving quick and straightforward results for possible nest coordinates was possible only when specifications were met. Therefore we show both methods we applied to find possible nest locations.

##### 4.1 Prepare kernel .csv files in Excel

At the storage location, open the corresponding .csv file of the kernel with a double right-click. Select column A with a left-click. Select the 'Data' tab. Choose the function 'Text to Columns'. The Text Conversion Wizard opens. Continue with a left-click. For delimiters, left-click on 'Comma', then left-click on 'Finish'. Delete the columns: SP_ID, id_2, and area. Save the file.

##### 4.2 Create pivot table from H95

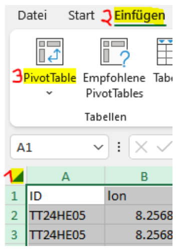


In the workbook, select the worksheet with the H95 data. Left-click the arrow (1) to select all cells. Left-click the 'Insert' tab (2). A window opens. The pivot chart should be created as a 'New Worksheet', so leave the existing selection as it is and confirm with a left-click on 'OK'. A new worksheet labeled 'Sheet1' appears. Change the worksheet name by double right-clicking on its label 'SheetX' and entering, for example, 'Pivot H95_TT24HE05'.

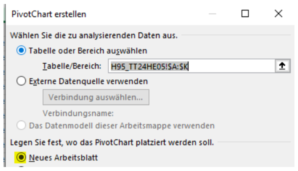


In the Pivot worksheet on the right, under PivotTable Fields, drag the line 'hour' with the cursor into the box under 'Columns' while holding the left-click. Drag the line 'hour' again into the box under 'Values'. It is automatically set to 'Sum of hour'. Left-click on this entry, and in the window that opens, left-click the bottom line: 'Value Field Settings'. In the subsequent window, under 'Summarize Value Field By', select the line: 'Count' and confirm with a left-click on 'OK'. Again, use the cursor on the PivotTable Fields window to drag 'date' into the box under 'Rows' while holding the left-click. This usually generates three entries. Left-click on the entry: 'Months' and in the window that opens, left-click on the line: 'Remove Field'. Repeat for the line: 'date'. Check if the days/hours follow each other seamlessly or if there are data gaps in the header (hours) or the dates.

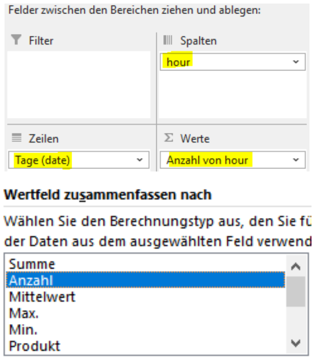


If needed, edit later in the Pivot H95 worksheet. To do this, first copy the data and insert it to the side, then add any missing time columns and date rows.

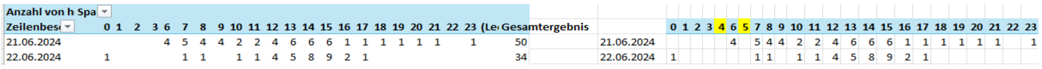

##### 4.3 Prepare nesting suspicion analysis sheet (example with core area)

In the Turtle dove project, an analysis is needed that displays the hours in a way that ensures eight consecutive hours can always be visible. For GPS data covering all hours of the day, additional columns will be required.

The analysis sheet is divided into three sections. On the left are the data of the kernel being analyzed. In the middle, these data are compared in percentage terms with the H95 data, which is considered 100%. The right section contains the H95 data. They are entered first.

C50 data on the left:
Columns: A - AI

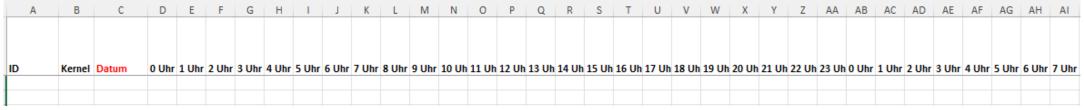


Analysis in the middle:
Columns: AJ - BP

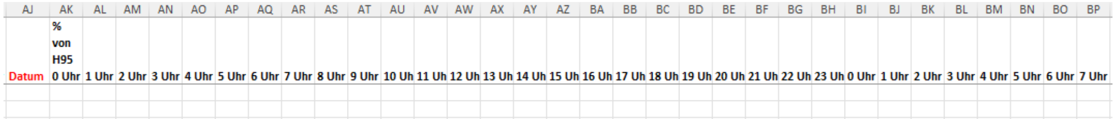


H95 data on the right:
Columns: BQ - CX

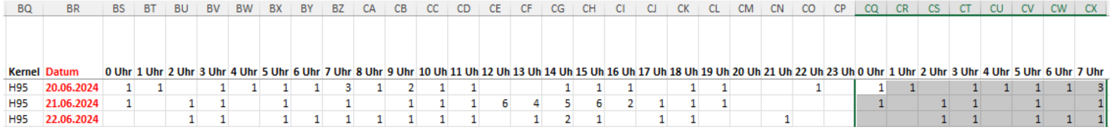


Column A = ID, Columns B, BQ = Kernel, Columns C, AJ, BR = Date, Columns D-AI, BS-CX = 0-23 and 0-7 hours, Column AK = % of H95 at 0 hours, Columns AL-BP = 1-23 and 0-7 hours. Create a new worksheet for this, naming it, for example, as Bv_C50 TTBB2401, and label the columns.

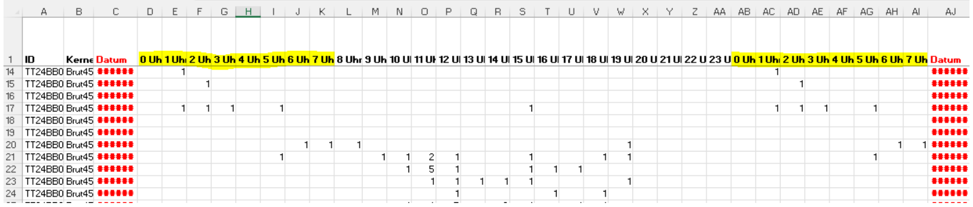

If the H95 data has no gaps in the header for hours or days, copy the entire data field with the date data and paste it into the worksheet Bv_C50 TTBB2401 starting from cell BR2. Then, copy the range from 0-7 hours to the last date row again and paste it starting from cell CQ2. Format the date in the desired format (DD.MM.YYYY). Copy the contents of the date column and paste them into the date column AJ. Create a pivot table from the C50 data worksheet in a similar manner and, if necessary, make adjustments. Insert these data starting from cell D2, then copy the data from 0-7 hours again and paste it starting from cell AB2. To be safe, save the file regularly.

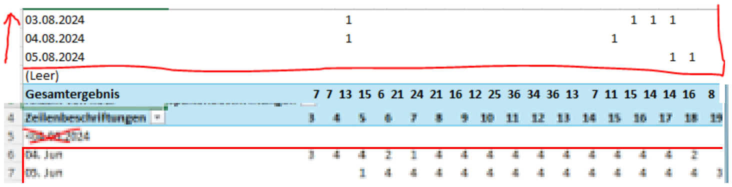

##### 4.4 Calculate percentage values of the chosen kernel (nestR kernel, core area, kernel 25) time values from H95 time values

In the first empty cell of the middle table, insert the formula adjusted for your table:

=IF(AND(firstTimeCellBackTable="", firstTimeCellFrontTable=""), "", IF(firstTimeCellFrontTable="", "--", (firstTimeCellFrontTable/firstTimeCellBackTable)*100))

For example, for times from 0-23 hours: =IF(AND(BS2="", D2=""), "", IF(D2="", "--", (D2/BS2)*100))

Explanation: If both BS2 and D2 are empty, nothing is displayed. If D2 is empty, "--" is shown. Otherwise, the percentage of D2 in relation to BS2 is calculated.

##### 4.5 Visualization of nesting suspicion windows

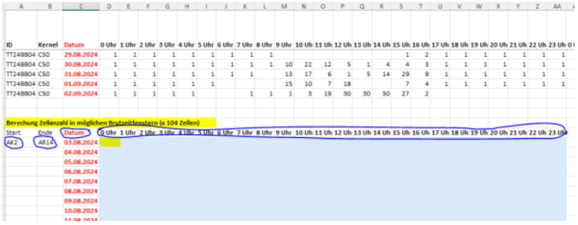

In turtle doves, usually two eggs are laid. They are incubated alternately by both parents, starting from the laying of the last egg. Incubation typically lasts 13-14 days. It is assumed that each bird incubates for at least about eight consecutive hours. The eggs are incubated for an average of approximately 80% of the time. The chicks are cared for by the parents for about 18-23 days after hatching. The female incubates at night, and the male during the day. There can be 1-3 broods per year. Therefore, sex plays a role in determining whether there is a nesting window in the middle block with percentage values when there is sufficient data.

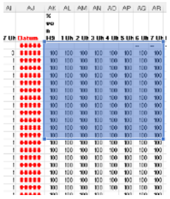

Our nesting window consists of 104 cells (8 consecutive hours over 13 consecutive days). We chose the shortest possible breeding time i.e., 13 days. To get an overview of possible nesting windows, a formula was developed. It provides an overview of which start cells for potential nesting windows are present in the middle percentage block and what the percentage of presence (here condition: >80%) is. It is recommended to apply this formula a few rows below the last row with data from the chosen kernel to be analyzed. Copy the data from the date column and paste it accordingly. Copy the time column from 0-23 hours and paste it into the adjacent cell. Insert the formula into the first free cell next to the first date and drag it to the following cells/rows using the small green square. Note: The end date of the GPS data must be considered.

| Datum | 0 Uhr | 1 Uhr | 2 Uhr | 3 Uhr | 4 Uhr | 5 Uhr | 6 Uhr | 7 Uhr | 8 Uhr |
|---|---|---|---|---|---|---|---|---|---|
| 18.08.2024 | 100 | 100 | 98 | 96 | 93 | 88 | 82 | | |
| 19.08.2024 | 100 | 99 | 97 | 95 | 92 | 87 | 81 | | |
| 20.08.2024 | 99 | 97 | 95 | 93 | 89 | 84 | | | |
| 21.08.2024 | 97 | 95 | 93 | 91 | 87 | 83 | | | |
| 22.08.2024 | 97 | 95 | 92 | 90 | 86 | 82 | | | |
| 23.08.2024 | 96 | 94 | 91 | 89 | 85 | 80 | | | |
| 24.08.2024 | Bleibt leer, da Folgezeit nicht für Brutverdacht ausreicht | | | | | | | | |
| 25.08.2024 | | | | | | | | | |
| 26.08.2024 | | | | | | | | | |
| 27.08.2024 | | | | | | | | | |
| 28.08.2024 | | | | | | | | | |
| 29.08.2024 | | | | | | | | | |
| 30.08.2024 | | | | | | | | | |
| 31.08.2024 | | | | | | | | | |
| 01.09.2024 | | | | | | | | | |
| 02.09.2024 | | | | | | | | | |
| 01.09.2024 | | | | | | | | | |
| 02.09.2024 | | | | | | | | | |

There can be no nesting windows in the date range that no longer allows for the end of a possible brood. For the TTs, these would be the last 12 days.

Formula:

=IF(SUM(StartCell:EndCell) / (COUNT(StartCell:EndCell) + COUNTIF(StartCell:EndCell, "--")) >= 80, SUM(StartCell:EndCell) / (COUNT(StartCell:EndCell) + COUNTIF(StartCell:EndCell, "--")), "")

For example, for times from 0-23 hours: =IF(SUM(AK2:AR14) / (COUNT(AK2:AR14) + COUNTIF(AK2:AR14, "--")) >= 80, SUM(AK2:AR14) / (COUNT(AK2:AR14) + COUNTIF(AK2:AR14, "--")), "")

Explanation: The Excel formula checks if the sum of the values in the range AK2:AR14 divided by the number of cells with numbers and the cells containing the value "--" is greater than or equal to 80. If this is the case, the average of these values is returned; otherwise, a blank value ("") is displayed. By double left-clicking on a cell in the analysis range, the corresponding nesting window in the middle percentage block is shown. If, as in the above example, there are many potential nesting windows, you can repeat the process with a smaller kernel (K25). Alternatively, it might be worthwhile to continue working directly in QGIS with the already categorized routes.

##### 4.6 Route analysis in QGIS

In the above example of TT24BB04A (female), it was shown that there are many potential nesting windows in C50 but none in K25. Therefore, it is sensible to conduct a route analysis.

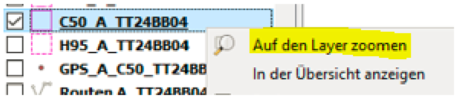


Open the QGIS project. Uncheck the layer with the background map (Google Satellite) and check the C50 layer (pink) — it consists of three areas here. Zoom to the layer by right-clicking on it in the layer list and selecting "Zoom to Layer"

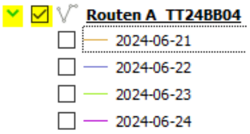

from the expanded options menu. Now check and expand the route layer of the individual. Then toggle the days on/off one by one.

In the example, it can be seen that in the left part of C50, there is no clustering of different-colored routes on the checked days. Several lines run through the middle C50 part, and a noticeable clustering is visible in the right C50 part. The further noticeable clustering above the middle C50 part can be ignored because it does not lie within C50, where we observed the multitude of nesting suspicion windows. A pattern as in the right C50 part could represent a nesting suspicion and should be verified.

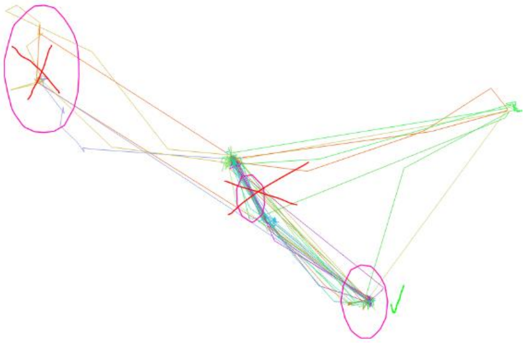

#### 4.7 Verification: create breeding point layer (core area, kernel 25)

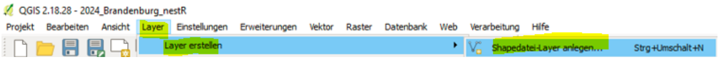


To do this, create a point layer. In the top menu bar, select the 'Layer' tab, then 'Create Layer', and choose 'Create Shapefile Layer' (all with a left-click).

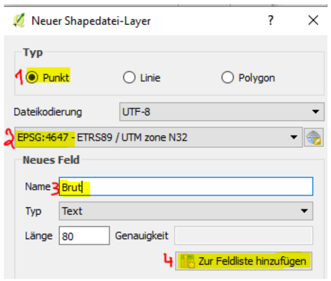


A window opens, and you enter: Point (1) under Type, 4647 (2) under CRS (Coordinate Reference System), and for Name, enter something like Breeding (3) (all with a left-click). Also, left-click on the option: 'Add to field list' (4) and confirm with a left-click on OK. A window opens and suggests a possible storage location. Check if it is the desired location; if not, browse to it and save the Esri shapefile under a name such as Breeding1. The layer appears in the layer panel.

Activate the layer with a left-click and enable the editing mode (pencil icon). Then, select 'Add Feature' (three points) with a left-click and place a point at the center of the crossed route lines. A window opens. Enter something like 1 in the Breeding field and confirm with a left-click on OK. Deactivate the editing mode for the layer and confirm with a left-click on Save when asked to save.

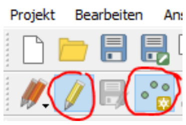


The coordinate data of the layer can be determined by saving the layer again in CRS: 4326, opening the attribute table, enabling the editing mode and the field calculator, creating a new field, and naming it something like: long. Then select 'Decimal number (real)' for Output field type and enter appropriate values (10, 6) for Output field length. In the expression box, enter $X and confirm with OK and save. Repeat this for the lat value (column name: lat) with the expression: $Y. Once you deactivate the editing mode and confirm the prompt to save changes, the generated columns remain. Save the coordinate data externally, and afterwards, for better clarity, remove the CRS 4326 layer in the layer panel.

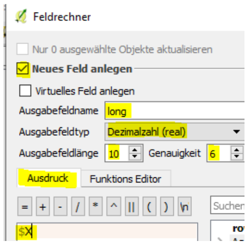


#### 4.8 Create buffer around point and intersect with GPS data

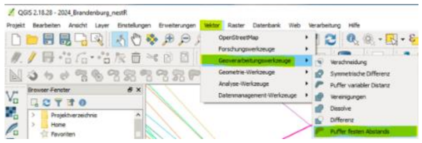

The point layer: Breeding should be active. In the menu bar, left-click: Vector ⟶ Geoprocessing Tools ⟶ Fixed distance buffer. A window opens. Select the Breeding layer under Input layer. Enter a distance that matches the GPS data scattering of your devices (+ possible safety margin - we used 45 m).

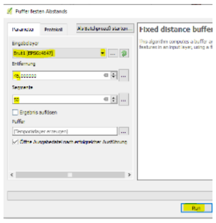

For segments, enter 50, to get a nice round buffer. Left-click on Run. A solid-colored temporary buffer appears in the layer list and on the map. Therefore, right-click on the buffer layer and save it as an Esri shapefile, for example: BreedingBuffer1 with: CRS 4647 at the desired location. Immediately delete the temporary buffer from the layer list. To do this, right-click on the layer. In the window that opens, select the option: Remove with a left-click and confirm with a left-click on OK. If necessary, change the appearance of the buffer area as described in 3.6.

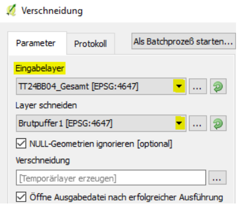


Then, Vector tab → Geoprocessing Tools → Intersection → Input layer: GPS data of the individual, Select BreedingBuffer1 as the intersect layer and confirm with a left-click on Run.

This creates a temporary intersection layer as described in 3.7. Continue as described in 3.8. If you have already conducted a nesting suspicion analysis for the individual, for example, with C50, it saves time to copy this worksheet, insert it again in the workbook, and rename it accordingly. Then delete the data from the left part and insert the resulting data (from the pivot table of the intersection with BreedingBuffer1) accordingly. Pay attention to dates and times: intersections often have gaps in days and times!

Note: The nesting suspicion analysis sheet can also be used for determining the sex in breeding birds like the turtle dove, where males incubate during the day and females incubate at night, if no blood sample is available or hasn't been analyzed yet. The turtle doves often don't stay in the breeding area when they're not incubating. They use the time to visit feeding habitats/perches, creating characteristic gaps in presence in the breeding area, which are easily recognizable as occurring during the nighttime for males and the daytime for females (see figure in 4.10).

Repeat with other results of breeding suspicion derived from the route analysis if nescessary.

##### 4.9 Using nestR to determine possible nest coordinates

We used nestR to confirm possible nest coordinates we determined using the breeding analysis sheets for core area and kernel 25.

To use nestR the free program JAGS needs to be installed on the computer. Download it using the link: https://sourceforge.net/projects/mcmc-jags/

| A | B | C | D |
|---|---|---|---|
| burst | date | long | lat |
| 2023_TT23SA01 | 2023-06-06 11:01:00 UTC | 12.14541 | 51.14458 |
| 2023_TT23SA01 | 2023-06-06 11:02:00 UTC | 12.14562 | 51.1444 |
| 2023_TT23SA01 | 2023-06-06 11:04:00 UTC | 12.14622 | 51.14433 |

Using nestR, the data of all individuals needs to be compiled into an Excel workbook, possibly named 'Total'. The required columns are: burst, date, long, and lat. Note: It is also possible to import workbooks for each individual, but this will take much more time, as the following procedure then needs to be repeated or each individual.

The burst column should contain the year and the bird ID, combined with an underscore. The date column contains the timestamp, while the long and lat columns contain the coordinates.

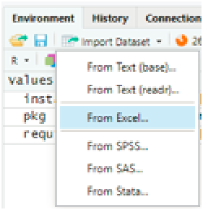


Import your workbook into the R Environment by left-clicking on 'Import Dataset' and then left-clicking on 'From Excel'. A window opens. Browse (1) to the destination where your data is stored. Use the down-pointing arrows to change the settings of date (2) to Date, long (3)

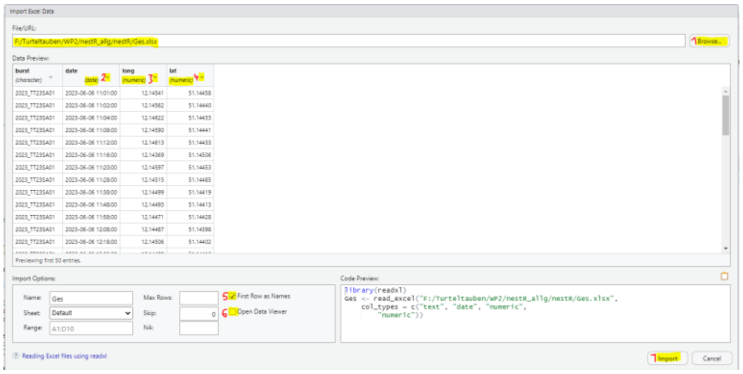

and lat (4) to Numeric each. Check if the box 'First Row as Names' is ticked (5) and untick 'Open Data Viewer' (6). Left-click on 'Import' (7).

Copy lines 77-124 of the R script 'Supp_info_BV.R' into the source pane, modify them to match your own project, and then run all lines by left-clicking on 'Run'. The Excel file with the possible nest coordinates will be saved again at the previously set storage location. Go there and open it. It will contain three sheets: 'Output_A1nests', 'A1nests', and 'A1visits'. You might find a multitude of possible nest coordinates for each individual. According to our experience, and with the possibility of turtle doves breeding 0-3 times per season, the first and second entry for each individual showed the true nest coordinates. In our data, the maximum number of detected breeding attempts was two. There might no results for some individuals, leading to the assumption that they did not breed in the given year.

| A | B | C | D |
|---|---|---|---|
| burst | loc_id | long | lat |
| 2023_TT23HE01 | 971 | 8.8697456 | 50.8618432 |
| 2023_TT23HE01 | 4053 | 8.870928 | 50.8615136 |
| 2023_TT23HE01 | 538 | 8.8800184 | 50.8591744 |
| 2023_TT23HE01 | 2519 | 8.8703632 | 50.8617216 |
| 2023_TT23HE01 | 1232 | 8.85156 | 50.8659872 |
| 2023_TT23HE01 | 2449 | 8.8591048 | 50.8619232 |

Proceed with creating .csv files with the possible nest coordinates for the individuals, using the first and second entries. Then, enter the new .csv files to your QGIS-project as described in 2.3-2.4. Afterwards proceed with 4.8 and then use 4.5-4.6 (if needed) to verify the coordinates.

##### 4.10 Check Potential Nesting Success

If nesting suspicion has been verified for an individual, you can additionally check if the nesting was potentially successful. In the case of turtle doves, the adult birds must care for their nestlings for at least another 18 days. Therefore, you can look at the nesting suspicion analysis sheet of the respective area to see if there are presences for at least another 18 days. For this, check the middle section (percentage comparison) to see if there are sufficient presences on the days following the nesting period (nesting period: pink) for the respective timeframe (nestling care: green), as shown in the illustrated example. For turtle doves, presences are significantly more pronounced in the first subsequent days than later, as the nestlings become increasingly independent and may leave the nest to be present in the surroundings. Here, a potential success was noted, as the adult bird was occasionally present in the nesting area for 20 days.

| AJ | AK | AL | AM | AN | AO | AP | AQ | AR | AS | AT | AU | AV | AW | AX | AY | AZ | BA | BB | BC | BD | BE | BF | BG | BH |
|---|---|---|---|---|---|---|---|---|---|---|---|---|---|---|---|---|---|---|---|---|---|---|---|---|
| Datum | % von H95 0 Uhr | 1 Uhr | 2 Uhr | 3 Uhr | 4 Uhr | 5 Uhr | 6 Uhr | 7 Uhr | 8 Uhr | 9 Uhr | 10 Uh | 11 Uh | 12 Uh | 13 Uh | 14 Uh | 15 Uh | 16 Uh | 17 Uh | 18 Uh | 19 Uh | 20 Uh | 21 Uh | 22 Uh | 23 Uh |
| 24.06.2024 | -- | 100 | 100 | 100 | -- | -- | 100 | -- | 100 | -- | -- | -- | -- | -- | -- | -- | -- | -- | -- | -- | -- | 100 | -- | -- |
| 25.06.2024 | -- | 100 | -- | 100 | 100 | 100 | 100 | 100 | -- | -- | -- | -- | -- | -- | -- | -- | -- | -- | -- | 9 | 75 | 100 | 100 | 100 |
| 26.06.2024 | -- | 100 | 100 | 100 | 100 | 100 | 100 | 100 | 100 | | -- | -- | -- | -- | -- | -- | -- | -- | -- | 50 | 100 | -- | 100 | 100 |
| 27.06.2024 | 100 | 100 | 100 | 100 | 100 | 100 | 100 | 100 | 100 | 100 | -- | -- | -- | -- | -- | -- | -- | -- | -- | -- | 100 | | -- | 100 |
| 28.06.2024 | 100 | 100 | 100 | -- | 100 | 100 | 100 | 100 | 100 | 100 | -- | -- | -- | -- | -- | -- | -- | -- | -- | 100 | -- | 100 | 100 | 100 |
| 29.06.2024 | -- | 100 | 100 | 100 | 100 | 100 | 100 | 100 | 100 | -- | -- | -- | -- | -- | -- | -- | -- | -- | -- | 100 | -- | 100 | 100 | 100 |
| 30.06.2024 | 100 | 100 | 100 | 100 | -- | -- | 100 | 100 | -- | 100 | | -- | -- | -- | -- | -- | -- | -- | -- | -- | 100 | 100 | 100 | -- |
| 01.07.2024 | 100 | 100 | 100 | 100 | 100 | 100 | 100 | 100 | -- | 100 | -- | -- | -- | -- | -- | -- | -- | -- | -- | 18 | 50 | -- | -- | 100 |
| 02.07.2024 | -- | -- | 100 | 100 | 100 | -- | | 100 | -- | 100 | -- | -- | -- | -- | -- | -- | -- | -- | -- | -- | 100 | -- | 100 | 100 |
| 03.07.2024 | 100 | 100 | -- | 100 | 100 | 100 | 100 | 100 | 100 | 100 | -- | -- | -- | -- | -- | -- | -- | -- | -- | -- | 100 | 100 | 100 | -- |
| 04.07.2024 | 100 | 100 | 100 | 100 | 100 | 100 | -- | 100 | 100 | 100 | -- | -- | -- | -- | -- | -- | -- | -- | -- | 100 | 100 | 100 | 100 | 100 |
| 05.07.2024 | 100 | 100 | 100 | | 100 | 100 | 100 | 100 | 100 | -- | -- | 100 | -- | | | -- | -- | -- | -- | -- | 100 | | 100 | 100 |
| 06.07.2024 | 100 | 100 | 100 | 100 | -- | 100 | 100 | 100 | 100 | 100 | 100 | -- | -- | -- | -- | -- | -- | -- | 14 | 50 | -- | 100 | 100 | 100 |
| 07.07.2024 | 100 | 100 | 100 | 100 | -- | -- | -- | 100 | 100 | -- | -- | -- | -- | -- | -- | -- | -- | -- | 14 | 100 | 100 | 100 | 100 | -- |
| 08.07.2024 | -- | 100 | 100 | 100 | 100 | -- | 100 | 100 | 25 | -- | -- | -- | -- | -- | -- | -- | -- | -- | -- | 33 | 67 | 50 | 100 | 100 |
| 09.07.2024 | 100 | 100 | 100 | 100 | 100 | 100 | 100 | 50 | 50 | -- | -- | -- | -- | -- | -- | -- | -- | -- | -- | 58 | 67 | 50 | -- | -- |
| 10.07.2024 | 100 | 100 | 100 | 100 | 100 | 100 | 100 | 100 | 100 | 50 | -- | -- | -- | -- | -- | -- | -- | 3 | 86 | 100 | 100 | 100 | | |
| 11.07.2024 | 100 | 100 | 100 | 100 | 100 | 100 | -- | 100 | -- | 100 | -- | -- | -- | -- | -- | -- | -- | 21 | 50 | 100 | 100 | 100 | 100 | -- |
| 12.07.2024 | 100 | | 100 | 100 | 100 | 100 | 100 | 100 | 100 | | -- | -- | -- | -- | -- | 31 | 100 | | | | | | | |
| 13.07.2024 | -- | 100 | | | | | | | -- | -- | -- | -- | -- | -- | 14 | 83 | 40 | -- | -- | 31 | 83 | 75 | -- | 100 |
| 14.07.2024 | -- | -- | 100 | 100 | 100 | -- | -- | -- | -- | -- | 8 | -- | -- | -- | -- | -- | -- | -- | 27 | -- | -- | -- | -- | -- |
| 15.07.2024 | -- | -- | -- | -- | -- | -- | -- | -- | -- | -- | -- | -- | -- | -- | -- | -- | 46 | -- | -- | -- | 9 | -- | -- | -- |
| 16.07.2024 | -- | -- | -- | -- | -- | -- | -- | -- | -- | -- | 100 | -- | -- | -- | -- | -- | 50 | 13 | -- | -- | -- | -- | -- | -- |
| 17.07.2024 | -- | -- | -- | -- | -- | -- | 100 | -- | -- | -- | -- | -- | -- | -- | -- | -- | -- | -- | -- | -- | -- | 100 | -- | -- |
| 18.07.2024 | -- | -- | -- | -- | -- | -- | -- | -- | -- | -- | -- | -- | -- | -- | -- | -- | -- | 38 | 20 | -- | -- | -- | -- | -- |
| 19.07.2024 | -- | -- | -- | -- | -- | -- | -- | -- | | -- | -- | -- | -- | -- | -- | -- | -- | 24 | 33 | 40 | -- | -- | -- | -- |
| 20.07.2024 | -- | -- | -- | -- | -- | -- | -- | -- | -- | -- | -- | -- | -- | -- | -- | -- | -- | 92 | 22 | -- | -- | -- | -- | -- |
| 21.07.2024 | -- | -- | -- | -- | -- | -- | -- | 100 | -- | -- | 25 | -- | 100 | 33 | 100 | 50 | -- | -- | -- | -- | -- | -- | -- | -- |
| 22.07.2024 | | | -- | -- | -- | -- | -- | -- | -- | 100 | -- | -- | -- | | | | | -- | -- | 27 | -- | | | |
| 23.07.2024 | -- | -- | | | | | | -- | -- | 5 | 5 | -- | -- | -- | -- | -- | -- | -- | -- | -- | -- | -- | -- | -- |
| 24.07.2024 | -- | -- | -- | -- | -- | -- | -- | -- | -- | -- | -- | -- | -- | -- | -- | 8 | 38 | -- | -- | | -- | -- | -- | -- |
| 25.07.2024 | -- | -- | -- | -- | -- | -- | -- | -- | | -- | -- | -- | -- | -- | 24 | 71 | -- | -- | -- | -- | -- | 100 | -- | -- |
| 26.07.2024 | -- | | -- | -- | -- | -- | -- | -- | 100 | 100 | -- | -- | -- | -- | | -- | | | | -- | | | | |

```r
# Section 3.1
# R script for kernel calculation in R -

# Save the project:
# Click the File tab -> choose "Save As" -> specify the desired location and project name -> click Save.


# List of required packages
required_packages <- c("adehabitatHR", "maptools", "raster", "rgdal", "geosphere", "terra")

# Install packages
installed_packages <- rownames(installed.packages())
for (pkg in required_packages) {
  if (!pkg %in% installed_packages) {
    install.packages(pkg)
  }
}

# Load packages
library(adehabitatHR)
library(maptools)
library(raster)
library(rgdal)
library(geosphere)
library(terra)
#__________________________________________
# Section 3.2
# Set the working directory to the folder where your data is located
setwd("Path:/Folder/Subfolder/Subsubfolder/...")

# Example:
setwd("F:/Turtledoves/Core_area_and_home_range/DataDownload_2024-09-02/2024_Turtledove_Data")

# Load the individual’s data file
# Example:
TT24HE05 <- read.table("TT24HE05.txt", header = TRUE, sep = "\t")
#__________________________________________
# Section 3.3
# The location data is converted into a suitable format
locs <- TT24HE05 # <- modify to your individual

# The following command lines do not need to be modified
# Prepare data for adehabitatHR by retrieving only the X and Y data
xyt <- subset(locs, select = c(lon, lat))

# Remove the ID
id <- subset(locs, select = ID)

# Create a SpatialPointsDataFrame for the adehabitatHR package
locs1 <- id
coordinates(locs1) <- xyt
class(locs1)

# kernel density estimation
kud <- kernelUD(locs1[,1], h = "href", kern = "epa", grid = 200, extent = 1)

# For the kernel: Home range (H = 95% of the data points (dp)), core area (C = 50% of the dp),
# and Kernel25 (K25 = 25% of the dp), the results of the kernel density estimation are
# converted to vectors.
homerange <- getverticeshr(kud, percent = 95)
core <- getverticeshr(kud, percent = 50)
kernel <- getverticeshr(kud, percent = 25)
#__________________________________________
# Section 3.4
# Visualization in R
plot(locs1)
plot(homerange,border=91, lwd=2, add=T) # plots homerange outline
```

```r
plot(core,border=4, lwd=2, lty = "solid", add=T) # plots corearea outline
plot(kernel,border=rainbow(10), lwd=2, lty = "solid", add=T) # plots kernel outline
#_________________________________________
# Section 3.5
# Creation of shapefiles
writePolyShape(homerange,"H95TT24HE05") # <- modify to your individual
writePolyShape(core,"C50TT24HE05")      # <- modify to your individual
writePolyShape(kernel,"K25TT24HE05")    # <- modify to your individual
#_________________________________________
# Section 4.9
# Use nestR to identify nesting attempts in turtle doves (TTs)

# Install required packages
install.packages("remotes")
install.packages("kableExtra")
install.packages("rpart")
install.packages("rpart.plot")
install.packages("forcats")
install.packages("DT")

# After installing JAGS on the computer, test if R recognizes it
library(rjags) # <- it worked

# Install nestR
remotes::install_github("picardis/nestR", build_vignettes = TRUE)

# Load required libraries
library(nestR)
library(readr)
library(dplyr)
library(writexl)
library(rpart)
library(rpart.plot)
library(forcats)
library(geosphere)
library(DT)
library(stringr)
library(lubridate)
library(ggplot2)
library(readxl)

# Import Excel data of all individuals compiled to one document into the environment, setting 'date' to Date and 'long' and 'lat' to Numeric
# First run with find_nests with discard_overlapping = FALSE
A1 <- find_nests(gps_data = Total,
                 sea_start = "03-01",   # Season start on 01.03.
                 sea_end = "09-30",     # Season end on 30.09.
                 nest_cycle = 36,       # 36 days = 18 days incubation + 18 days chick rearing
                 buffer = 45,           # corresponds to the buffer used in QGIS for nesting suspicion analysis
                 min_pts = 2,           # loose settings, as in Picardi
                 min_d_fix = 5,         # loose settings, as in Picardi
                 min_consec = 18,       # during nesting each partner must be at the nest for at least 18 days (with at least 8 consecutive hours)
                 min_top_att = 1,       # loose settings, as in Picardi <- percentage value
                 min_days_att = 1,      # loose settings, as in Picardi <- percentage value
                 discard_overlapping = FALSE) # uncertain which arguments differentiate a real nest from a similarly frequented location

# Save file as Excel
write_xlsx(A1,"A1.xlsx")
```
